# Differential modulation of the cellular and humoral immune responses in *Drosophila* is mediated by the endosomal ARF1-Asrij axis

**DOI:** 10.1101/056796

**Authors:** Rohan J. Khadilkar, D.R. Chetan, Arghyashree RoyChowdhury Sinha, Srivathsa S. Magadi, Vani Kulkarni, Maneesha S Inamdar

## Abstract

How multicellular organisms maintain immune homeostasis across various organs and cell types is an outstanding question in immune biology and cell signaling. In Drosophila, blood cells (hemocytes) respond to local and systemic cues to mount an immune response. While endosomal regulation of *Drosophila* hematopoiesis is reported, the role of endosomal proteins in cellular and humoral immunity is not well-studied. Here we demonstrate a functional role for endosomal proteins in immune homeostasis. We show that the ubiquitous trafficking protein ADP Ribosylation Factor 1 (ARF1) and the hemocyte-specific endosomal regulator Asrij differentially regulate humoral immunity. ARF1 and Asrij mutants show reduced survival and lifespan upon infection, indicating perturbed immune homeostasis. The ARF1-Asrij axis suppresses the Toll pathway anti-microbial peptides (AMPs) by regulating ubiquitination of the inhibitor Cactus. The Imd pathway is inversely regulated-while ARF1 suppresses AMPs, Asrij is essential for AMP production. Several immune mutants have reduced Asrij expression, suggesting that Asrij co-ordinates with these pathways to regulate the immune response. Our study highlights the role of endosomal proteins in modulating the immune response by maintaining the balance of AMP production. Similar mechanisms can now be tested in mammalian hematopoiesis and immunity.

## Introduction

A cascade of appropriate responses to infection or injury is dynamically regulated to co-ordinate the immune response. However, mechanistic details of how the immune organs and molecules they produce communicate, are poorly understood. In the open circulatory system of *Drosophila*, hemocytes carry out phagocytosis and melanization whereas the humoral immune response is mediated by the fat body and gut. Plasmatocytes, which form a majority of the hemocyte population, phagocytose invading pathogens, crystal cells melanize and restrict pathogens to the affected area and lamellocytes encapsulate and neutralize large objects such as parasites^1^. A serine protease cascade in crystal cells activates prophenoloxidase (ProPO), which then catalyzes the conversion of phenols to quinones that then polymerize to form melanin^2^. A Toll pathway-dependent protease inhibitor Serpin27A produced by the fat body is required to limit melanization to crystal cells^3,4^. Thus mechanisms of transport and uptake are essential to regulate systemic communication and melanization. Larvae and adults deficient in Serpin27A or the ProPo mutant *Black cells* show spontaneous melanization yet are immune compromised^5^.

Humoral immunity is primarily governed by the Toll and Imd (Immune deficiency) pathways, which regulate anti-microbial peptide (AMP) production. Fungi or Gram positive bacteria induce the Toll pathway, which causes activation of the NF-KB factor Dif and production of AMPs such as Drosomycin, Metchnikowin and Defensin. Infection by Gram negative bacteria causes activation of the Rel homology and IkappaB homology domain protein Relish, through the Imd pathway and leads to the production of Diptericin, Attacin, Cecropin and Drosocin^6^. Additionally, immune pathways have also been shown in tissue-specific contexts. The JAK/STAT pathway regulates gut-mediated defense mechanisms by controlling intestinal stem cell proliferation^7,8^ and is also essential for the production of humoral factors like thio-ester proteins and turandot molecules in response to septic injury^9^. The receptor tyrosine kinase Pvr also plays an important role in regulating the Imd pathway. *Drosophila* JNK activates the expression of Pvr ligands, Pvf2 and Pvf3 which bind Pvr and lead to the activation of the Pvr/ERK pathway which negatively regulates the JNK and NF-κB arm of the Imd pathway^10^.

AMPs are primarily produced by the fat body, the equivalent of the mammalian liver, and secreted into circulation to reach target tissues^11^. The systemic response by the fat body is mainly governed by the Toll and Imd pathways^12^. An intriguing question is the mode of communication between the hemocyte and fat body-mediated immune response. Few studies have recently shown that the hemocytes contribute towards fat body mediated immune responses. Psidin is a lysosomal protein required for degradation of bacteria in hemocytes and simultaneously required for Defensin production in the fat body^13^ and Spaetzle has been shown to have a paracrine effect on the fat body mediated immune response^14,15^. Normal physiological levels of AMPs are altered in response to infections but can also be affected by genetic mutations, thus perturbing homeostasis similar to an infection-induced condition^16–19^.

Just as in hematopoiesis, a complex network of molecular interactions is essential to regulate immune function and maintain homeostasis. Hence understanding organismal regulation of these processes is a major challenge. Elucidating mechanisms that can integrate the varying inputs received at the cellular level and orchestrate the outcome will be key to generating tools that allow control of the system. Several recent studies highlight the importance of unique signal generation and regulation in cellular organelles, such as endosomes, in a variety of cell types^20–26^.

Endosomal regulation is a potent mechanism to integrate and modulate signals in *Drosophila* hematopoiesis^27,28^. We recently showed that the GTP bound form of the ubiquitous endosomal protein ADP Ribosylation Factor 1 (ARF1-GTP), interacts with and regulates the hemocyte-specific endosomal protein Asrij to integrate and modulate multiple signaling pathways required to maintain blood cell homeostasis, including the JAK/STAT and Notch pathways and Pvr and Insulin signaling^26,29^. Depletion of ARF1 or Asrij leads to increased circulating and differential hemocyte counts. As the primary role of *Drosophila* hemocytes is to mount an immune response like their mammalian blood cell counterparts, and because both ARF1 and Asrij have conserved functions, we analyzed their requirement in the cellular and humoral immune response of *Drosophila*. We show that crystal cell number in ARF1 and Asrij mutants correlates with the extent of melanization. Further, perturbation of the ARF1-Asrij axis results in aberrant AMP production and compromises the response to bacterial infection. ARF1 and Asrij have similar effects on the Toll pathway but regulate Imd pathway AMPs inversely. Thus, we demonstrate that regulation by endosomal proteins allows differential response to signals to achieve immune homeostasis.

## Results

### Depletion of ubiquitous (ARF1) or hemocyte-specific (Asrij) endosomal proteins does not affect phagocytosis

Phagocytosis is a cellular immune response brought about by plasmatocytes, a differentiated hemocyte type. Depletion of either ARF1 or Asrij results in increased plasmatocyte numbers^26,29^. To analyze whether there is any alteration in the phagocytic ability of the mutant hemocytes we incubated the ARF1 knockdown or *asrij* null mutant larval circulating hemocytes with India ink dye particles^30^. Our analysis shows that phagocytosis is unaltered in both mutant genotypes as compared to controls (Fig. S1A-D). Thus the ARF1-Asrij axis is not essential for phagocytosis.

### Increased crystal cell number upon ARF1 or Asrij depletion co-relates with increased melanization and phenoloxidase activity

Crystal cells usually attach to the larval cuticle and mechanical injury or infection triggers melanization, leading to blackening of crystal cells^31^. Larvae depleted of ARF1 using a hemocyte driver (*e33cGAL4*) as well as *asrij* null mutant larvae, have increased crystal cells^26 29^. To test whether the mutant crystal cells were functional we subjected them to a melanization assay (see methods). Both mutant genotypes showed increase in melanized cells upon heat activation (about 1.5 fold for ARF1 knockdown and 2 fold for *asrij* mutant), as compared to controls (Fig. 1 A-B, F] and 2A-B, F). The depleted phenotypes could be rescued to near control levels by over-expressing the respective protein in larvae using *e33cGal4* (Fig. 1E-F and 2E-F). Overexpression of ARF1 or Asrij in wild type animals resulted in reduced crystal cell number corresponding to reduced differentiation seen in these larvae as reported earlier^26,29^ (Fig. 1C-D, F and 2C-D, F). Notably, the size and intensity of melanized spots was higher in the *asrij* null mutant compared to controls and Asrij overexpressing larvae (Figure 2A-F). This suggested increase in crystal cell number may be accompanied by increased function.

**Figure 1:**
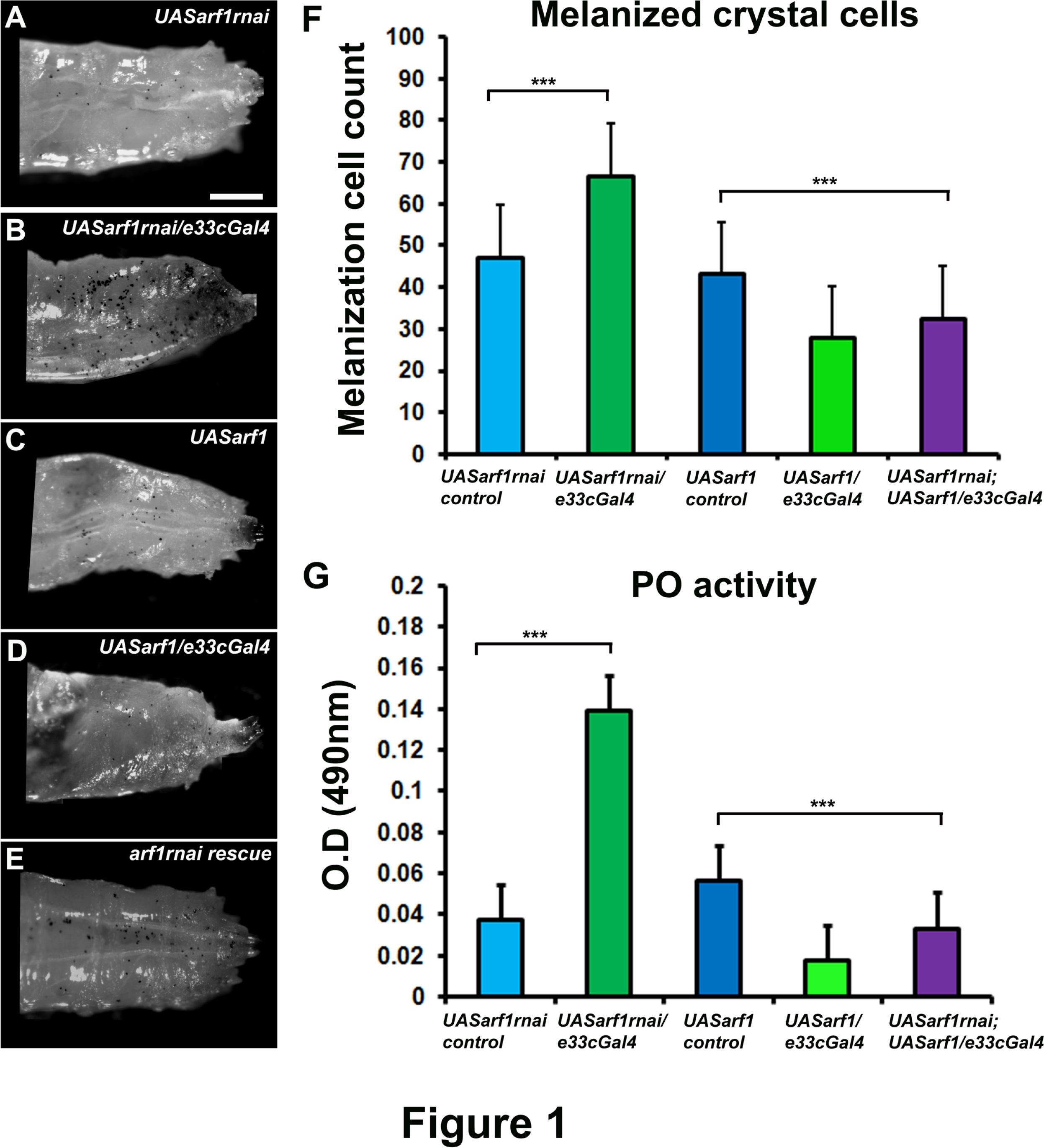
ARF1 regulates crystal cell-mediated melanization and phenoloxidase activity. (A-E) Photomicrographs showing posterior region of third instar larvae of specific genotypes (A) *UASarf1rnai* (B) *UASarf1rnai/e33cGal4* (C) *UASarf1* (D) *UASarf1/e33cGal4* (E) *UASarf1rnai/UASarfrnai; UASarf1/e33cGal4* that were heated at 60°C for 10 min to visualize the melanization response. (F) Melanized crystal cells were quantitated and represented graphically. (G) Graph representing phenoloxidase activity in the hemolymph of the indicated genotypes (detected as absorbance at OD490nm) after conversion of L-3, 4-dihydroxyphenylalanine. Scale Bar: (A-E) 100μm. Number of larvae analyzed per genotype (n=10). Error bars show standard deviation of mean. P-value: ^***^ indicates P<0.001.

**Figure 2:**
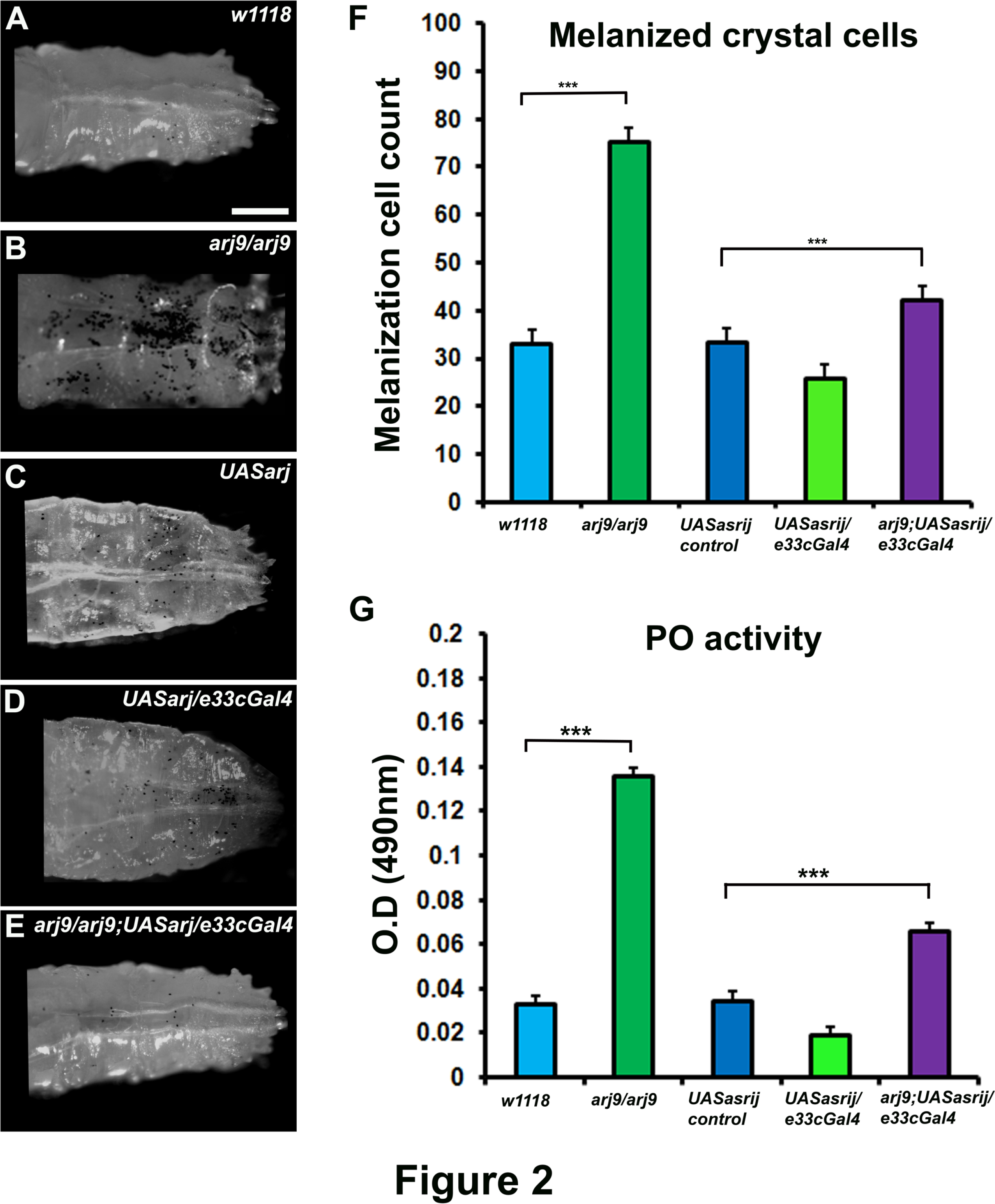
Asrij regulates crystal cell-mediated melanization and phenoloxidase activity. (A-E) Photomicrographs showing posterior region of third instar larvae of specific genotypes (A) *w1118* (B) *arj^9^/arj^9^* (C) *UASarj* (D) *UASarj/e33cGal4* (E) *arj^9^/arj^9^; UASarj/e33cGal4* that were heated at 60°C for 10 min to visualize the melanization response. (F) Melanized crystal cells were quantitated and represented graphically. (G) Graph representing phenoloxidase activity in the hemolymph of the indicated genotypes (detected as absorbance at OD490nm) after conversion of L-3, 4-dihydroxyphenylalanine.Scale Bar: (A-E) 100μm. Number of larvae analyzed per genotype (n=10). Error bars show standard deviation of mean. P-value: ^***^ indicates P<0.001.

To test crystal cell function we assayed for phenoloxidase (PO) activity. Melanin biosynthesis is brought about by PO, which catalyzes the oxidation of phenols to quinones, which subsequently polymerize into melanin. A protease cascade cleaves the inactive zymogen proPO (PPO) to generate active PO. Both ARF1 knockdown and *asrij* null mutant larvae showed similarly increased PO activity (4 fold above control), which was completely restored by ARF1 overexpression but only partially restored (2 fold above control) upon Asrij over-expression (Fig. 1G and 2G). Interestingly, either ARF1 or Asrij over-expression in a wild type background reduced PO activity significantly, as expected from the reduced crystal cell number.

### ARF1 and Asrij cooperatively regulate expression of Toll pathway AMPs

Activation of the Toll pathway results in production of a repertoire of AMPs mainly, Drosomycin in response to fungal infection and Metchnikowin and Defensin in response to infection by Gram positive bacteria^32–35^. Under normal conditions these peptides are expressed at very low levels. However, by quantitative polymerase chain reaction-based analysis of reverse transcribed RNA (qRT-PCR) from uninfected adult flies, we found that transcript levels of *drosomycin* and *metchnikowin* are highly upregulated in ARF1 knockdown flies (*e33cGAL4*>*UAS:ARF1-RNAi*) (5.5 fold and 3.5 fold respectively) compared to the GAL4 control whereas *defensin* expression is not significantly altered (0.5 fold) (Fig. 3A). These data indicate that ARF1 depletion leads to activation of Toll pathway target genes even in the absence of infection. Asrij mediates ARF1 function by interacting with ARF1-GTP during Drosophila hematopoiesis. Further ARF1 regulates Asrij expression and stability. Hence we also checked the effect of Asrij depletion on AMP transcript levels. In the *asrij* null mutant we saw modest increase in *drosomycin* and *metchnikowin* transcript levels as compared to *w1118* control whereas *defensin* expression was slightly reduced (Fig. 3B). These data indicate that Asrij depletion also leads to differential activation of Toll pathway target genes. However a significant difference was the substantially greater effect of ARF1 depletion on *drosomycin* and *metchnikowin* transcript levels than that of Asrij depletion. This suggests that ARF1 has a wider role in regulating AMPs of the Toll pathway.

**Figure 3:**
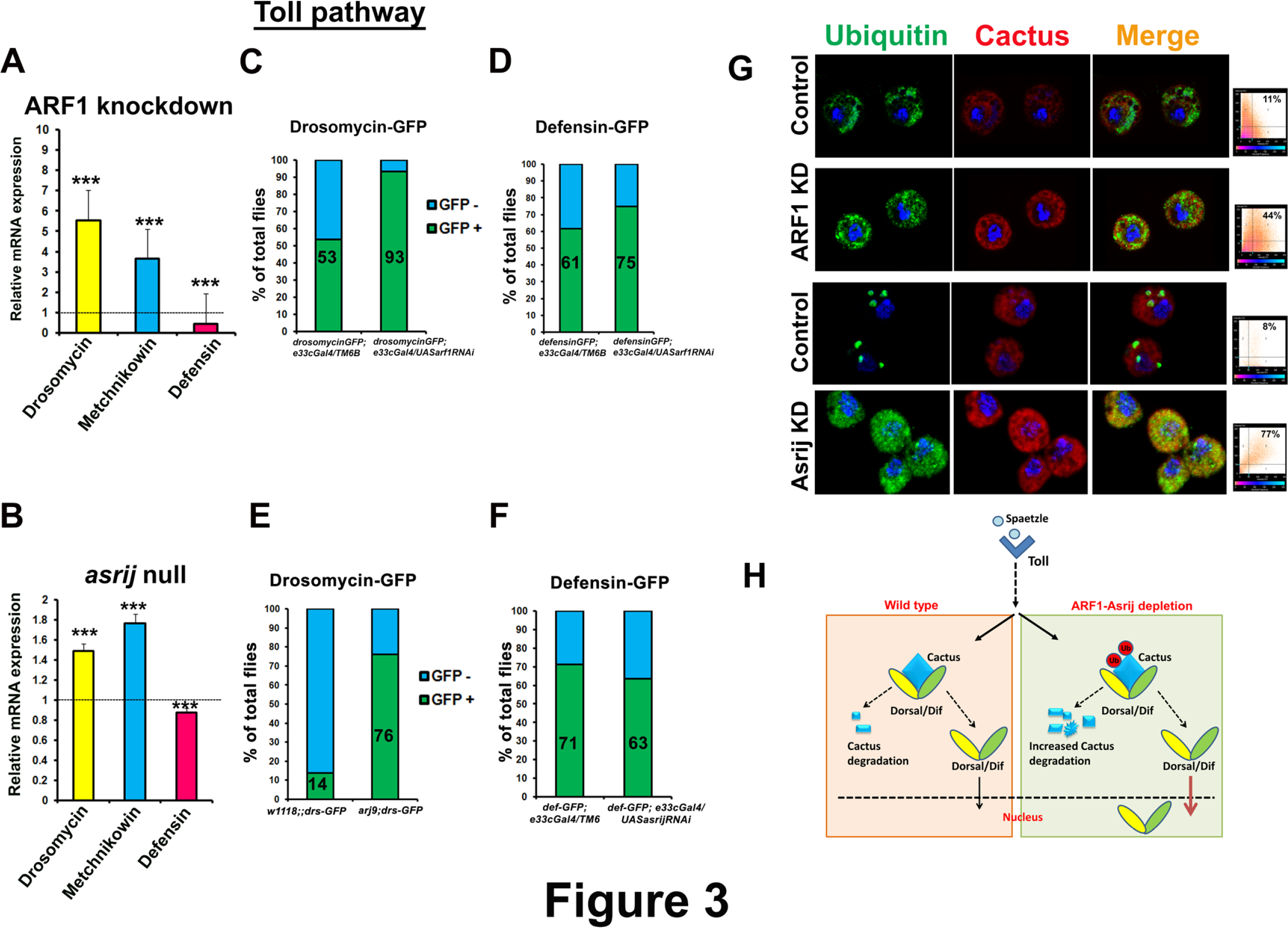
ARF1 and Asrij negatively regulate Toll pathway-mediated immune response. (A-B) Quantification of Toll pathway-governed antimicrobial peptide expression by qRT-PCR analysis shows that *Drosomycin* and *Metchnikowin* are upregulated and *Defensin* is downregulated in *ARF1* knockdown (A) and *asrij* null mutant (B) larvae. (C-F) Quantitation for the total percentile of flies expressing the Toll pathway reporters - Drosomycin-GFP and Defensin-GFP in flies with *e33cGal4*-mediated *ARF1* knockdown (C,D) or *asrij* null (E) or *e33cGal4*-mediated *asrij* knockdown (F) respectively. (G) Images showing increased colocalization of Cactus and Ubiquitin in ARF1 knockdown or *asrij* null circulating larval hemocytes as compared to respective controls, also indicated by adjacent co-localization plots (H) Model indicating the suggested role of the ARF1-Asrij endocytic axis in regulating the Toll pathway. Scale Bar: (G) 10μm. Error bars show standard error of mean. P-value: ^***^ indicates P<0.001.

We also performed *in vivo* reporter assays to test the activation status of representative Toll pathway AMPs using transgenic flies that would express GFP under the control of an AMP promoter upon infection. Introduction of Drosomycin-GFP or Defensin-GFP reporter in ARF1 knockdown or in *asrij* null or *asrij* knockdown background respectively was assayed after infection with *B. subtilis*. Quantitation of the GFP+ flies showed that ARF1 knockdown, resulted in significantly more GFP positive flies for Drosomycin (93% compared to 53% in GAL4 control) and a smaller increase in Defensin-GFP (75% compared to 61% in GAL4 control) (Fig. 3C,D). Similarly *asrij* depletion resulted in greatly increased Drosomycin-GFP expressing flies (76% compared to 14% in control) with a small reduction in Defensin-GFP flies (63% compared to 71% in control) (Fig. 3E, F). Thus our *in vivo* reporter analysis is in agreement with the mRNA expression analysis.

### The ARF1-Asrij axis suppresses Toll pathway AMP production by stabilizing Cactus

Activation of the Toll pathway leads to nuclear translocation of the transcription factors Dorsal and Dif. In the absence of the Toll ligand Spaetzle, Dorsal and Dif are bound by Cactus, a negative regulator of the Toll pathway, inhibiting their activity and nuclear localization^36^. Receptor activation leads to phosphorylation of Cactus followed by its ubiquitination and degradation, thus releasing Dorsal/Dif to translocate to the nucleus for target AMP gene activation. Since Toll pathway AMPs were upregulated in both Asrij and ARF1 depleted conditions we probed the status of ubiquitinated Cactus in these conditions. Depletion of ARF1 or Asrij led to increased co-localization of Cactus with Ubiquitin (Fig. 3G) indicating that it is increasingly targeted for degradation when the ARF1-Asrij axis is perturbed. Thus the ARF1-Asrij axis may maintain homeostatic conditions of Toll signaling by stabilizing Cactus and preventing aberrant AMP production (Fig. 3H).

### Differential effect of ARF1 and Asrij on Imd pathway AMPs

Infection with Gram negative bacteria brings about nuclear localization of the NF-κB transcription factor Relish, thus activating its main target AMP, Diptericin^32,33,35^. The other Imd pathway target genes include *attacin A1, cecropin A1* and *drosocin*. We analyzed the effect of ARF1 depletion on the transcript levels of Imd pathway targets and found that they were constitutively upregulated in ARF1 knockdown flies. *attacin*, *drosocinand* and *diptericin* transcripts were highly upregulated (~18 fold, ~15 fold and ~22 fold) whereas *cecropin* levels showed modest increase (1.25 fold) as compared to the parental control (Fig. 4A). Upon *in vivo* AMP-GFP reporter analysis after infection with *E. coli*, ARF1 knockdown showed a dramatic up-regulation of the percent of GFP positive flies for Attacin, Cecropin, Drosocin and Diptericin (87.5%, 88.8%, 100% and 84.21% respectively) as compared to Gal4 control (68.75%, 77.7%, 54.5% and 63.6% respectively) (Fig 4C-F).

**Figure 4:**
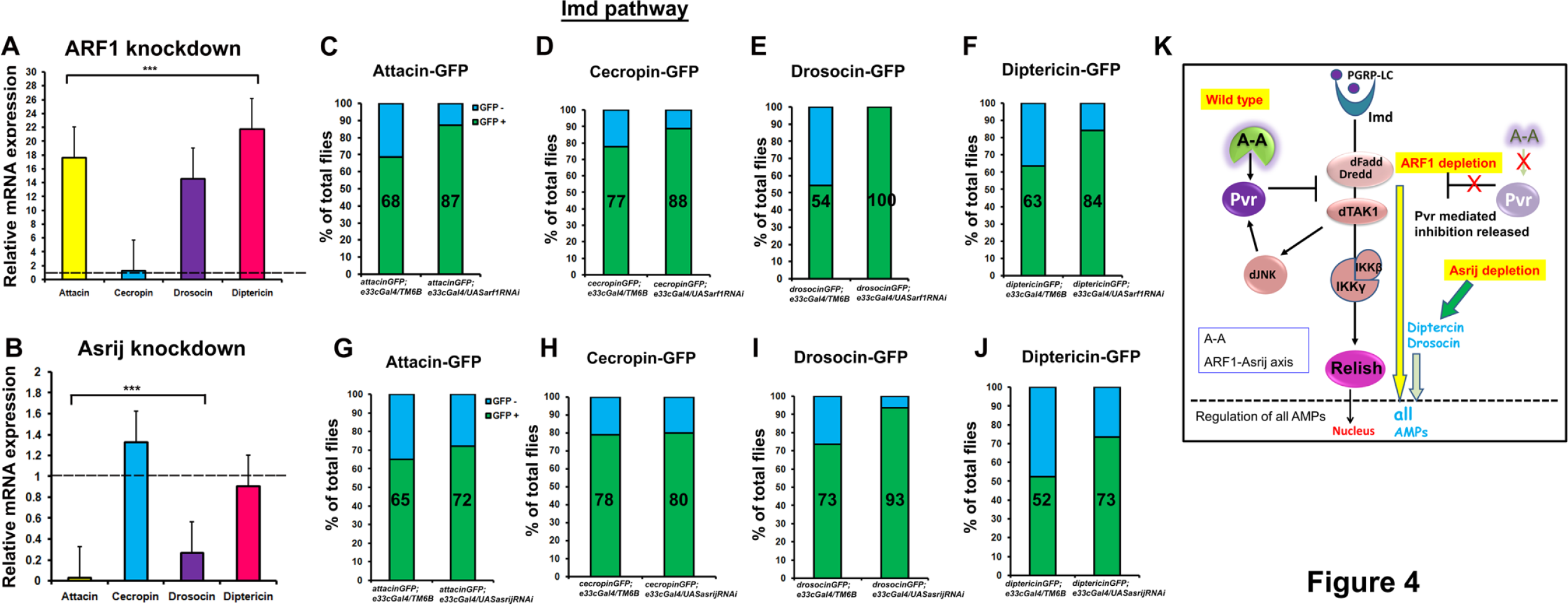
ARF1 and Asrij differentially regulate the Imd pathway. (A-B) Quantification of Imd pathway-governed antimicrobial peptide expression by qRT-PCR analysis shows that *Attacin*, *Drosocin* and *Diptericin* are highly up-regulated whereas *Cecropin* levels are unaffected in *e33cGal4*-mediated *ARF1* knockdown flies (A). *Cecropin* levels are upregulated and *Attacin*, *Drosocin*, Diptericin levels are down-regulated in *e33cGal4*-mediated Asrij knockdown flies (B). (C-J) Quantitation of the total percentile of flies expressing the Imd pathway reporters - Attacin-GFP, Cecropin-GFP, Drosocin-GFP and Diptericin-GFP in flies with *e33cGal4* mediated *ARF1* knockdown (C-F) or *asrij* knockdown (G-J) respectively. (K) Model indicating the suggested role of the ARF1-Asrij endocytic axis in regulating the Imd pathway. Error bars show standard error of mean. P-value: ^***^ indicates P<0.001.

In contrast to the effect of *ARF1* depletion, *Asrij* null mutants showed no significant change in expression of *diptericin* transcripts, which is a standard indicator of Imd activation (Fig 4B). However, while levels of *attacin* and *drosocin* were reduced, *cecropin* transcript levels were marginally affected (1.4 fold) and comparable to ARF1 knockdown flies. In *vivo* AMP-GFP reporter analysis upon E. *coli* infection for the Imd pathway governed AMPs showed no significant change in GFP+ flies for Attacin and Cecropin (72.22%, 80% in knockdown compared to 65%, 78.94% in controls respectively) whereas Drosocin and Diptericin GFP+ flies were significantly higher (93.75% and 73.68% in knockdown compared to 73.68% and 52.63% in controls respectively) (Fig 4G-J).

These data show that both ARF1 and Asrij have major non-overlapping roles in regulating Imd pathway AMP expression. ARF1 is a generic negative regulator of the Imd pathway, as its depletion leads to heightened levels of all the target AMPs. While there is only a small change in *diptericin* or *cecropin* transcript levels upon Asrij depletion, Diptericin peptide levels are higher indicating Asrij normally checks Diptericin levels, possibly post-transcriptionally. While Asrij positively regulates *attacin* and *drosocin* transcript expression, the effect on Attacin AMP levels was not significant. However there was a dramatic increase in Drosocin-GFP flies. This indicates that Asrij shows differential/ discriminatory effects on AMP production (Fig. 4K). Thus co-ordinated and complementary regulation of Imd targets by the ARF1-Asrij axis is essential to maintain immune homeostasis.

### Survival and lifespan of Asrij or ARF1-depleted flies is compromised upon acute bacterial infection

Impaired AMP production upon infection can lead to reduced survival due to an inability to combat the bacterial infection. Loss-of-function mutations in the Toll and Imd pathway effectors such as Dif and Relish^37^ lead to reduced ability to combat infections. Earlier studies show that Asrij is epistatic to ARF1 and depends on ARF1 for its stability^26^. While they both similarly regulate the Toll pathway, differential regulation of the Imd pathway suggests complex control on AMP production. Hence we tested the effect of Asrij or ARF1 depletion on the ability of flies to combat infection and survive.

Upon infection with *B. subtilis*, while 80% control flies continued to survive after 48 hrs with a gradual reduction in number over subsequent days, mutant numbers rapidly declined after 48 hrs and the mutant population perished 3-4 days earlier than controls. This resulted in a steep decrease in % survival upon infection as compared to the *Gal4* and *w1118* controls respectively (Fig 5A, C). Thus the increased production of the Toll pathway AMP Drosomycin in the absence of ARF1 or Asrij does not protect lifespan upon Gram positive bacterial infection.

**Figure 5:**
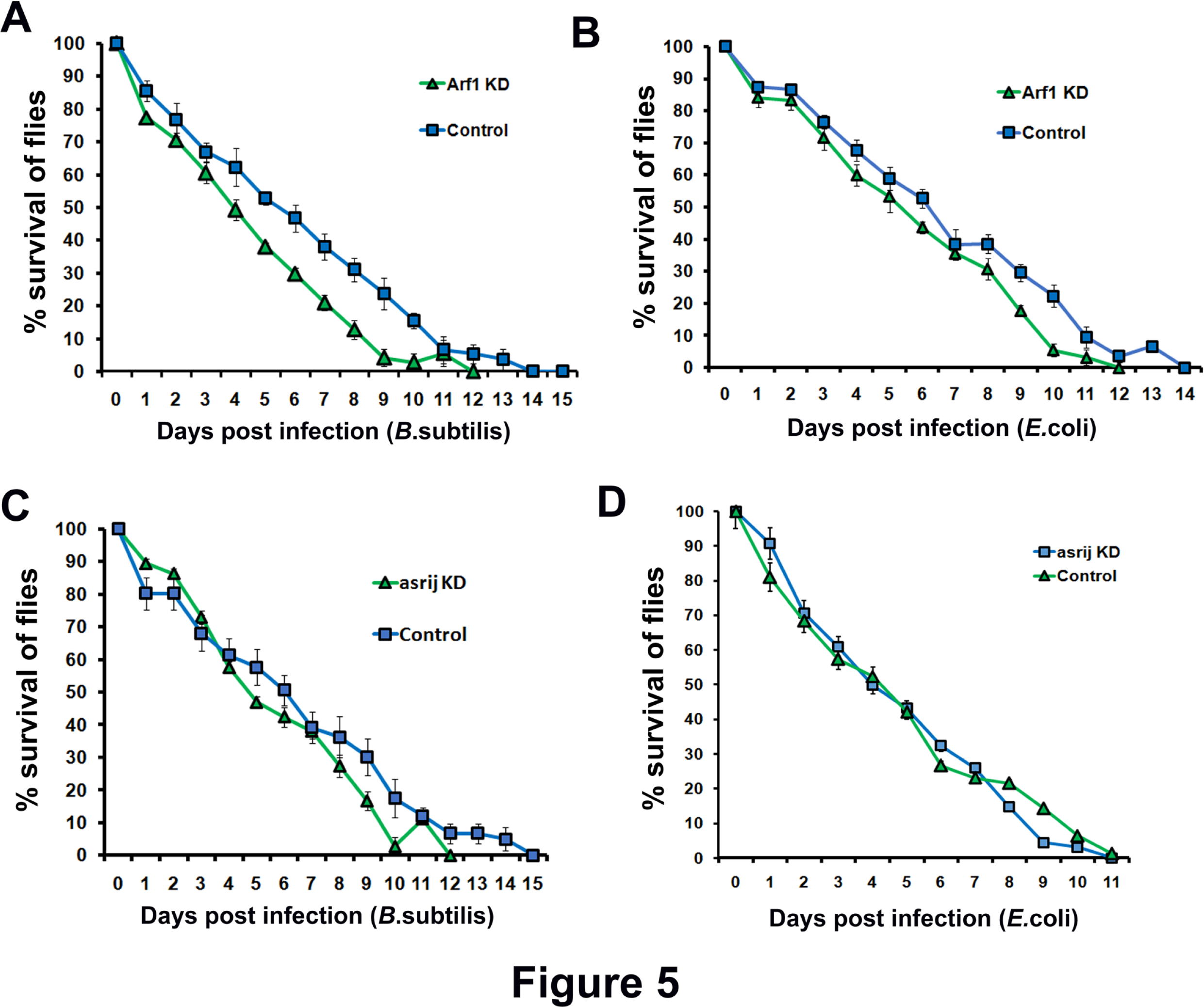
ARF1 and Asrij knockdown flies show compromised survival upon infection. (AD) Survival curves showing that *e33cGal4*-mediated *ARF1* or *asrij* knockdown flies show a reduced survival ability as compared to its control upon infection with either *B*.subtilis (A, C) or *E*.coli (B, D). At least 100 flies were tested per genotype over at least three independent experiments.

While *ARF1* depletion results in increased production of all Imd pathway AMPs, *asrij* depletion caused limited and differential activation of Imd pathway AMPs. Upon infection with *E.coli*, flies depleted of ARF1 or Asrij, both showed compromised survival (Fig.5B,D). While *asrij* depletion made flies more susceptible than controls, the effect of *ARF1* knockdown was less dramatic. However all mutant genotypes perished before controls, indicating that increased AMP production does not confer the ability to combat infection.

### Asrij expression is downregulated upon Gram negative bacterial infection

Immune response in normal animals requires AMP upregulation to combat infection^6,8,38^. ARF1 and Asrij have opposite effects on the Imd pathway, suggesting that Asrij acts independent of ARF1 in Imd pathway regulation. However, increased AMP levels in ARF1 knockdown do not provide additional ability to combat infection. Additionally, Asrij is downstream of ARF1 and reduces AMP levels. Therefore Asrij expression levels positively correlate with Imd pathway target AMP transcript levels. Hence we checked the status of Asrij expression upon immune challenge. Wild type flies infected with *E.coli* showed reduced *asrij* transcript levels (Fig. 6A). This indicates that *asrij* levels co-relate with Imd pathway activation.

**Figure 6:**
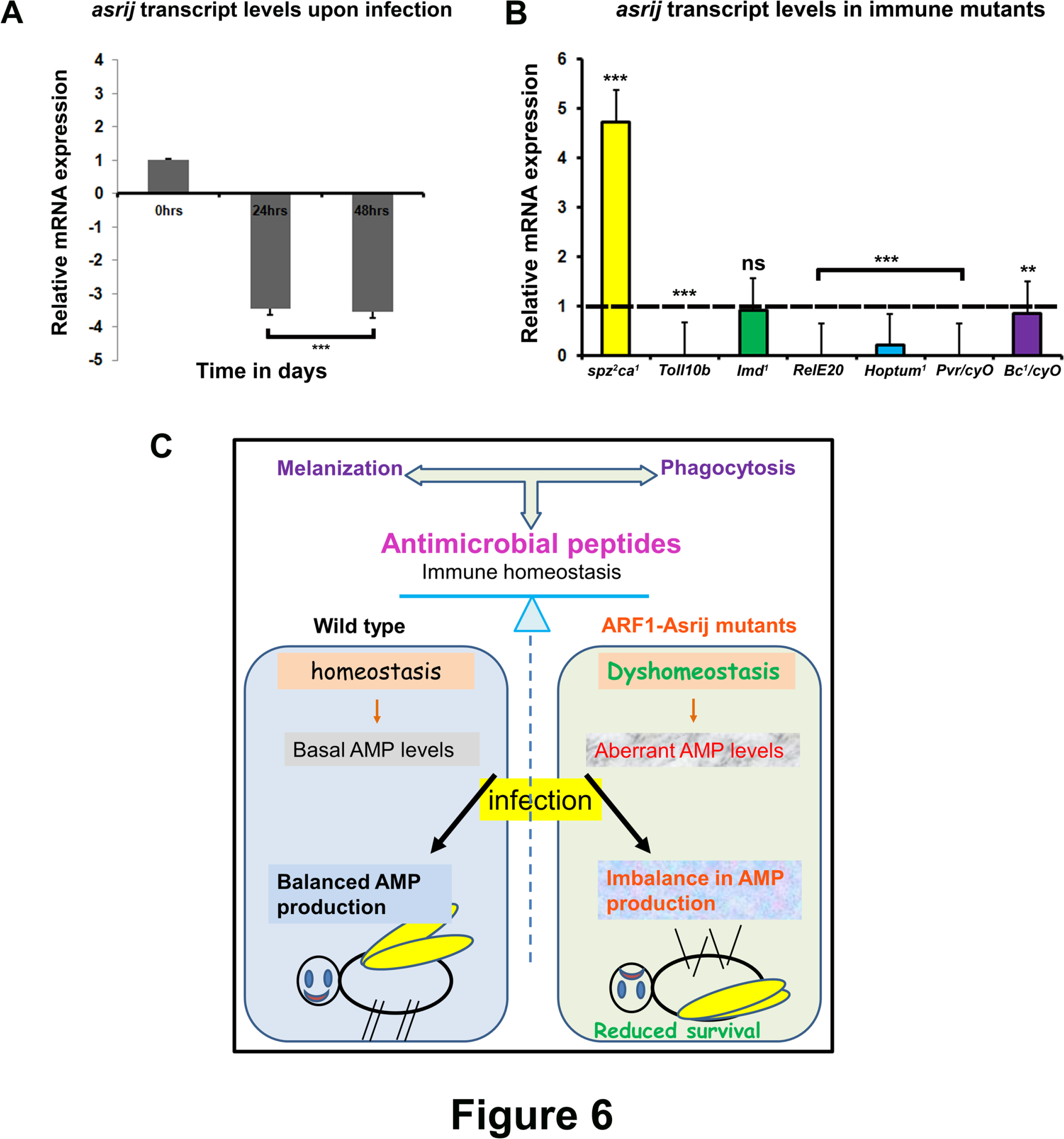
Asrij expression is modulated upon infection and in immune mutants. (A-B) qRT-PCR analysis of adult flies showing that *asrij* mRNA levels are down-regulated 24 and 48 hours post infection (A) and that *asrij* transcript levels vary among immune mutants (B). (C) Model illustrating loss of immune homeostasis in ARF1-Asrij depleted conditions leading to compromised survival of the flies.

### Asrij levels are differentially regulated in immune pathway mutants

Asrij levels are downregulated upon bacterial infection and survival is compromised in Asrij depleted conditions indicating an important role for Asrij in mounting an immune response. Since pathway activation and attenuation are both important for maintaining homeostasis, it is likely that Asrij expression is in turn regulated by components of the Toll or Imd pathways as well as other pathways like Jak/Stat and Pvr. To test this we assayed *asrij* transcript levels in a range of immune pathway mutants which either inactivate or activate the pathway. We found that *asrij* levels are upregulated in Toll pathway mutants like *spf^rm7^* whereas they are downregulated in Toll pathway gain-of-function mutants like *Toll^10b^* (Fig. 6B). This is in agreement with increased ubiquitination of cactus, suggesting a requirement for the ARF1-Asrij axis in regulating Toll pathway activity possibly via a feed-forward loop.

*Asrij* levels are downregulated in Imd pathway mutants like *Rel^E20^* (Imd effector) suggesting that *asrij* may be a target of *relish*. Also mutants of the other pathways like *Hoptum^1^* (Jak gain of function) and *Pvr* (Pvr/CyO) show reduced Asrij levels (Fig. 6B). There was no significant downregulation of *asrij* levels in *Imd* and *Black cells* mutant. This suggests that *asrij* is differentially regulated by downstream effectors of the Toll and Imd immune pathways. While the Toll pathway suppresses *asrij* expression, Imd pathway promotes *asrij* expression. Taken together with our data that Asrij levels are reduced upon E. coli infection this suggests that Asrij expression may be dependent on Imd pathway activation.

## Discussion

A balanced cellular and humoral immune response is essential to achieve and maintain immune homeostasis^19,39,40^. In *Drosophila*, aberrant hematopoiesis and impaired hemocyte function can both affect the ability to fight infection and maintain immune homeostasis. Endosomal proteins are known to regulate *Drosophila* hematopoiesis^26,29^. Here, we show an essential function for endosomal proteins in regulating immunity.

Altered hemocyte number and distribution as a result of defective hematopoiesis, can also lead to immune phenotypes like increased melanization or phagocytosis. Perturbation of normal levels of ARF1 or Asrij leads to aberrant hematopoiesis, affecting the circulating hemocyte number^26,29^. We show that this in turn leads to an impaired cellular immune response. In addition, ARF1 and Asrij have a direct role in humoral immunity by regulating AMP gene expression. Thus we propose a model for the role of the ARF1-Asrij axis in maintaining immune homeostasis (Figure 6C) that can be used for testing additional players in the process.

ARF1 is involved in clathrin coat assembly and endocytosis^41^ and has a critical role in membrane bending and scission^42^. In this context it is also intriguing that ARF1, like Asrij, does not seem to have an essential role in phagocytosis. This suggests that hemocytes could be involved in additional mechanisms beyond phagocytosis in order to combat an infection.

Both ARF1 and Asrij control hemocyte proliferation as their individual depletion leads to an increase in the total and differential hemocyte counts. Also, both mutants have higher crystal cell numbers due to over-activation of Notch as a result of endocytic entrapment^26,29^. This suggests that increased melanization accompanied by increase in phenoloxidase activity upon ARF1 or Asrij depletion is a consequence of aberrant hematopoiesis not likely due to a cellular requirement in regulating the melanization response. Constitutive activation of the Toll pathway or impaired Jak/Stat or Imd pathway signaling in various mutants also leads to the formation of melanotic masses^33^. Thus the phenotypes seen on Asrij or ARF1 depletion could be an outcome of loss of control over the immune regulatory pathways.

Regulation of many signaling pathways, including the immune regulatory pathways takes place at the endosomes^43–46^. For example, endocytic proteins Mop and Hrs co-localize with the Toll receptor at endosomes and function upstream of MyD88 and Pelle, thus indicating that Toll signaling is regulated by endocytosis^45^. Our study shows that loss of function of the ARF1-Asrij axis leads to an upregulation of some AMP targets of the Toll pathway. Upon depletion of ARF1-Asrij endosomal axis we find increased ubiquitination of Cactus, a negative regulator of the Toll pathway. Thus the endosomal axis may control the sorting and thereby degradation of Cactus. This could explain the significant increase in Toll pathway reporter expression such as Drosomycin-GFP. Interestingly the effect of ARF1 depletion on the Toll pathway is more pronounced than that of Asrij depletion. This is not surprising as ARF1 is a ubiquitous and essential trafficking molecule that regulates a variety of signals. This suggests that ARF1 is likely to be involved with additional steps of the Toll pathway and may also interact with multiple regulators of AMP expression.

ARF1 and Asrij show complementary effects on IMD pathway target AMPs. While ARF1 causes repression of transcription and production of IMD pathway AMPs, Asrij has a discriminatory role. Asrij seems to promote transcription of AttacinA and Drosocin, whereas it represses Cecropin. However in terms of AMP production only Drosocin and Diptericin are affected, but not to the extent of ARF1. Thus ARF1 causes strong generic suppression of the Imd pathway while the role of Asrij could be to fine tune this effect. Mass spectrometric analysis of purified protein complexes indicates that ARF1 and Imd interact^47^ (*Drosophila* Protein Interaction Mapping Project, https://interfly.med.harvard.edu). Hence it is very likely that ARF1 regulates Imd pathway activation at the endosomes. Whether this interaction involves Asrij or not remains to be tested and will give insight into modes of differential activation of immune pathways. Our analysis shows that Asrij is the effector for endosomal regulation of the humoral immune response by ARF1 and provides specialized tissue-specific and finer control over AMP regulation. This is in agreement with earlier data showing that Asrij acts downstream of ARF1^26^. Since ARF1 is expressed in the fat body^26^ it could communicate with the hemocyte-specific molecule, Asrij, to mediate immune cross talk.

As we see reduced Asrij expression in Toll and Jak/Stat pathway mutants such as *Rel^E20^* and *Hoptum^1^*, it is likely that these effectors also regulate Asrij, setting up a feedback mechanism to modulate the immune response. Endosomal regulation of NICD transport and activity and activation of Stat92e are both mediated by the ARF1-Asrij axis. Further, ARF1 regulates Pvr and Insulin signaling^26,29^. Asrij levels are also downregulated in the *Pvr* mutant. ARF1 is known to act downstream of Pvr in HSC maintenance and Asrij acts in concert with ARF1^26^. Hence it is likely that the ARF1-Asrij axis regulates trafficking of the Pvr receptor. This suggests active modulation of signal activity and outcome at endosomes could be orchestrated by ARF1 and Asrij.

AMP transcript level changes upon ARF1 or Asrij depletion also correspond to reporter-AMP levels seen after infection. This suggests that though ARF1 is known to have a role in secretion, mutants do not have an AMP secretion defect. Hence aberrant regulation of immune pathways on perturbation of the ARF1-Asrij axis is most likely due to perturbed endosomal regulation.

ARF1 has a ubiquitous function in the endosomal machinery^41^ and is well-positioned to regulate the interface between metabolism, hematopoiesis and immunity in order to achieve homeostasis. Along with Asrij and other tissue-specific modulators it can actively modulate the metabolic and immune status in *Drosophila*. In this context, it is interesting to note that Asrij is a target of MEF2^48^, which is required for the immune-metabolic switch *in vivo^49^*. Thus Asrij could bring tissue specificity to ARF1 action, for example, by modulating insulin signaling in the hematopoietic system.

It is likely that in Asrij or ARF1 mutants, the differentiated hemocytes mount a cellular immune response and perish as in the case of wild type flies where immunosenescence sets in with age and the ability of hemocytes to combat infection declines^50^. Since their hematopoietic stem cell pool is exhausted, they may fail to replenish the blood cell population, thus compromising the ability to combat infections. Alternatively, mechanisms that downregulate the inflammatory responses and prevent sustained activation^39,51^ may be inefficient when the trafficking machinery is perturbed. This could result in constitutive upregulation thus compromising immune homeostasis^51,52^.

In summary, we show that in addition to its requirement in hematopoiesis, the ARF1-Asrij axis can differentially regulate humoral immunity in *Drosophila*, most likely by virtue of its endosomal function. ARF1 and Asrij bring about differential endocytic modulation of immune pathways and their depletion leads to aberrant pathway activity and an immune imbalance. In humans loss of function mutations in molecules involved in vesicular machinery like Amphyphysin I in which clathrin coated vesicle formation is affected leads to autoimmune disorders like Paraneoplastic stiff-person syndrome^53^. Synaptotagmin, involved in vesicle docking and fusion to the plasma membrane acts as an antigenic protein and leads to an autoimmune disorder called Lambert-Eaton myasthenic syndrome^54^. Mutations in endosomal molecules like Rab27A, β subunit of AP3, SNARE also lead to immune diseases like Griscelli and Hermansky-Pudlak syndrome^55,56^. Mutants of both ARF1 and Asrij are likely to have drastic effects on the immune system. Asrij has been associated with inflammatory conditions such as arthritis, thyroiditis, endotheliitis and tonsillitis (http://www.malacards.org/card/tonsillitis?search=OCIAD1), whereas the ARF family is associated with a wide variety of diseases. ARF1 has been shown to be involved in mast cell degranulation and IgE mediated anaphylaxis response^57^. Generation and analysis of vertebrate models for these genes such as knockout and transgenic mice will provide tools to understand their function in human immunity.

## Methods

**Drosophila Stocks:** All fly stocks were maintained at standard rearing conditions. Respective UAS or Gal4 parent strains or *w1118* (*asrij* null mutant) were used as controls. Tissue specific Gal4 promoter line was used to drive the expression of UAS responder genes. Following fly lines were used: *arj^9^/arj^9^*, *UAS-asrij* (Kulkarni, Khadilkar et al., 2011), *relish^E20^*, Black cells *(Bc^1^/CyO)*, *Hoptum^1^*, *Imd^1^* (NCBS stock centre), Pvr/Cyo (Pernille Rorth, Denmark), Gal4 driver lines e33cGAL4 (Kathryn Anderson, NY, USA), *UASarf1rnai* (VDRC), GFP reporter flies for Toll and Imd pathway were a kind gift from David Ferrandon, France.

### Phagocytosis assay

Primary hemocyte cultures were prepared as described earlier^58^. Briefly, third instar larvae were surface sterilized, and hemolymph was collected by puncturing the integument using dissection forceps into 150 μl of 1X PBS (Phosphate Buffer Saline) in 35-mm coverslip-bottom dishes and incubated with India Ink (HIMEDIA, India) for 10 min followed by 5 washes with PBS. After 1 hour hemocyte preparations were fixed with 2.5% paraformaldehyde for 20 min and imaged. Phagocytosis of India ink particles by primary hemocytes was assessed using Zeiss LSM 510meta.

### Crystal cell melanization assay

Crystal cells are characterized by crystalline inclusions that contain the zymogen ProPO and can be visualized due to specific blackening upon heating larvae at 60 °C for 10 min^59^. Third instar wandering stage larvae heat treated to visualize crystal cells and imaged using SZX12 stereo zoom microscope (Olympus). Melanised crystal cell were counted from three posterior abdominal segments of at least 20 larvae per genotype. Error bars represent the standard deviation. P-values were calculated using one way ANOVA.

### Prophenol oxidase activity assay

For the measurement of PO activity by dot blots, 5 μl hemolymph was applied to a filter paper pre-soaked with 20 mM L-DOPA (L-3, 4-dihydroxyphenylalanine-Cat. No. D9628, SIGMA) in phosphate buffer pH 6.6, incubated for 20 minutes at 37 °C and heated in a microwave till the paper was dried completely^60^. The melanised black hemolymph spots correlate with PO activity in hemolymph and were imaged under an Olympus SZX12 stereo zoom microscope.

For photometric measurement of PO activity, 10 μl hemolymph from each strain was pooled on ice by quickly bleeding 3-5 larvae and withdrawing 6 μl hemolymph aliquots of mixed hemolymph were activated at 25 °C for 10 minutes, then 40 μl L-DOPA was added and optical density (OD) measured from 0 to 30 minutes at 490 nm with a Vmax^™^ Kinetic Microplate Reader (*BIO-RAD*). Activation of PO was measured as the relative change in absorbance (A_490_). Experiments were repeated at least three times with biological and technical replicates.

### RNA extraction and Quantitative Real Time PCR

*Drosophila* larvae or adults were collected in TRIzol (TRIzol^®^ Reagent, Cat. No. 15596-026, Invitrogen), homogenized for 30-60 seconds and centrifuged. The supernatant was processed for RNA extraction according to the manufacturer’s protocol. RNA was quantified by spectrophotometry and quality was analyzed by agarose gel electrophoresis. Quantitative RT-PCR (qRT-PCR) was performed using SYBR green chemistry in a Rotor Gene 3000 (Corbett Life Science) and analyzed with the accompanying software. Primer sequences used for RT-PCR and qRT-PCR are provided in supplementary information (Table S2).

### Infection and survival assay

Briefly, prior to infection, adult flies of appropriate genotype were starved for 2 hr, then transferred into vials containing filter paper hydrated with 5 % sucrose solution mixed with concentrated Ampicillin resistant *E.coli* (A_600_ = 1; concentrated to contain ~10 CFU/ml) or Gram positive bacteria (*B.subtilis*) A600= 5-10) on cornmeal food. Following incubation at 25 °C for 24 hr, flies were processed for RNA extraction or examined for reporter-GFP expression. For survival assay flies were challenged with bacteria by oral feeding starved adult flies starved for 48 hours then transferred to fresh corn-meal food vials containing fresh filter paper disks inoculated with bacterial cultures. The percentage of survivors was then calculated for each experiment and plotted as a survival curve. (N=50) for each genotype. Reporter-GFP expressing flies were imaged on an SZX12 stereo zoom microscope (Olympus) and processed uniformly for brightness/contrast using Adobe Photoshop CS3.

### Statistical analysis

Student’s t-test with unequal variances has been used for statistical analysis. P-values are as indicated in the graphs.

## Acknowledgements

This work was funded by a grant to MSI from the Department of Science and Technology, Government of India and by intramural funds from the Jawaharlal Nehru Centre for Advanced Scientific Research.

